# Ohana: detecting selection in multiple populations by modelling ancestral admixture components

**DOI:** 10.1101/546408

**Authors:** Jade Yu Cheng, Fernando Racimo, Rasmus Nielsen

**Affiliations:** Centre for GeoGenetics, Natural History Museum of Denmark, University of Copenhagen, Oster Voldgade 5-7, Copenhagen 1350 Denmark; Departments of Integrative Biology and Statistics, University of California, Berkeley, Berkeley, CA 94720, USA

**Keywords:** Positive selection, admixture, population structure, human evolution, selective sweeps

## Abstract

One of the most powerful and commonly used methods for detecting local adaptation in the genome is the identification of extreme allele frequency differences between populations. In this paper, we present a new maximum likelihood method for finding regions under positive selection. The method is based on a Gaussian approximation to allele frequency changes and it incorporates admixture between populations. The method can analyze multiple populations simultaneously and retains power to detect selection signatures specific to ancestry components that are not representative of any extant populations. We evaluate the method using simulated data and compare it to related methods based on summary statistics. We also apply it to human genomic data and identify loci with extreme genetic differentiation between major geographic groups. Many of the genes identified are previously known selected loci relating to hair pigmentation and morphology, skin and eye pigmentation. We also identify new candidate regions, including various selected loci in the Native American component of admixed Mexican-Americans. These involve diverse biological functions, like immunity, fat distribution, food intake, vision and hair development.

## Introduction

The emergence of population genomic data has facilitated fine-scale detection of regions under recent positive selection in humans and other species. There are multiple different methods for carrying out such selection scans. Some of these methods rely on patterns of long-range linkage-disequilibrium (Sabeti *et al.*, 2007;Voight *et al.*, 2006), one of the characteristic genomic footprints left by a selective sweep (Kim and Stephan, 2002;Kim and Nielsen, 2004;McVean, 2007). However, this pattern fades rapidly over time, and these methods are, consequently, best suited for detecting very recent selective sweeps from *de novo* mutations. Other methods, based on distortions in the allele frequency spectrum caused by positive selection, can allow for the detection of more ancient events, but are generally only applicable one population at a time (Tajima, 1989;Fu and Li, 1993;Fay and Wu, 2000;Nielsen, 2005;Huber *et al.*, 2016;DeGiorgio *et al.*, 2016).

A different class of methods for detecting selection analyses patterns of allele frequency differentiation between populations. These methods proceed, for example, by computing Wright’s fixation index (*F*_*ST*_) locally across different regions of a genome(Beaumont and Nichols, 1996;Akey *et al.*, 2002;Beaumont and Balding, 2004). The basic idea is that regions that have experienced episodes of positive selection will display frequency differences between populations that are stronger than what would be expected under pure genetic drift. Population differentiation methods can detect more ancient selective events than linkage disequilibrium-based methods (Sabeti *et al.*, 2006), and are sensitive to different types of positive selection events, including sweeps from a *de novo* mutation, sweeps from standing variation, incomplete sweeps, and adaptive introgression (Yi *et al.*, 2010;Bonhomme *et al.*, 2010;Fumagalli *et al.*, 2015;Racimo *et al.*, 2016). Recent methods have allowed researchers to detect excess local differentiation on particular branches of a 3-population tree (Yi *et al.*, 2010;Racimo, 2016), a 4-population tree (Cheng *et al.*, 2017) or an abitrarily large tree (Librado and Orlando, 2018), albeit without modeling post-split admixture events.

A generalization of these methods was, 2015;Coop *et al.*, 2010). Their method can handle an arbitrary number of populations and detects positive selection as a genomically local distortions from a genome-wide covariance matrix, which is used as a neutral baseline. Similar methods have used hierarchical Bayesian models (Foll and Gaggiotti, 2008;Foll *et al.*, 2014) or principal component analysis (Duforet-Frebourg *et al.*, 2015) to model patterns of population differentiation to identify local distortions across the genome. Another method ((Fariello *et al.*, 2013)) extended single-locus differentiation-based methods to the analysis of haplotype differentiation. More recently, Mathieson *et al.* (2015) developed an admixture-aware selection test based on a linear model and applied it to human data. The analysis took advantage of the fact that present-day European populations could be modeled as a mixture of three highly differentiated ancestral components. Regions of the genome that exhibited strong deviations from the genome-wide mixture proportions were therefore strong candidates for positive selection. Finally, Refoyo-Martinez *et al.* (2018) developed a method to test for selection on an admixture graph, which represents the history of divergence and admixture events among populations. Although useful for detecting selection in the presence of admixture, it still requires the user to specify which individuals belong to which populations, and to infer the graph in advance.

Here, we introduce a new selection detection framework that can explicitly model admixture and detect selection from populations of admixed ancestries. It can simultaneously compare arbitrarily many populations and ancestry components and is encoded in a flexible framework for testing selection on a specific lineage or set of lineages. The method allows the user to identify signals of positive selection via population differentiation, without relying on self-reported ancestry or admixture correction to group individuals into populations. The method can also determine if a selective event is specific to a particular population or shared among different populations.

Unlike previous methods, we fully take advantage of admixed populations, and we do not require the user to *a priori* categorize samples into populations, or to correct allele frequencies to account for recent admixture. Thus, the selection scan does not rely on user-supplied sample labels or ancestry compositions. The methods identifies positive selection by searching for loci showing distortions in the population covariance matrix, relative to the genome-wide baseline. It provides a flexible framework to specifically test for selection on individual components or sets of components. This functionality allows researchers to accommodate specific evolutionary scenarios into the range of testable hypotheses, including local adaptation, adaptive introgression, and convergent selection. The method first co-estimates the population structure of the input panel and the allele frequencies of the ancestral admixture compomnents through an unsupervised learning process (Cheng *et al.*, 2017), before testing for selection on the ancestral components themselves. Researchers can also use the method to examine estimated population structure and visualize trees connecting the ancestral components using plotting functionalities provided by our software package, Ohana, as part of the analysis pipeline.

## Methods

### Basic model

The new method is based on the Ohana inference framework (Cheng *et al.*, 2017), which works with both genotype calls and genotype likelihoods. In brief, the classical Structure model (Pritchard *et al.*, 2000) is used to infer allele frequencies, ancestry components, and admixture proportions using maximum likelihood (ML). Then a covariance matrix among components is inferred using a multivariate Gaussian distribution while enforcing constraints imposed by the assumption of a tree structure. This system is underdetermined (see e.g., Felsenstein (1985)), i.e. multiple covariance matrices induce the same probability distribution on the allele frequencies. To circumvent this issue, we root the tree in one of the ancestry components. This corresponds to conditioning on the allele frequencies in one of the components when calculating the joint distribution of allele frequencies in the other components. This idea is similar to Felsenstein’s restricted maximum likelihood approach (Felsenstein, 1985). We emphasize that the rooting is arbitrary but that it does not imply any assumptions about this component actually being ancestral.

Through ML estimation we obtain the covariance matrix Ω′, which has size (*K* − 1)*×*(*K* − 1) and a joint density shown in Eq. 1, where *f*_*kj*_ is the estimated allele frequency for ancestry component *k* at SNP *j* and *µ*_*j*_ is the sample allele frequency for SNP *j*, obtained either by counting alleles in the case of called genotypes or by EM estimation in the case of genotype likelihoods (Cheng *et al.*, 2017).

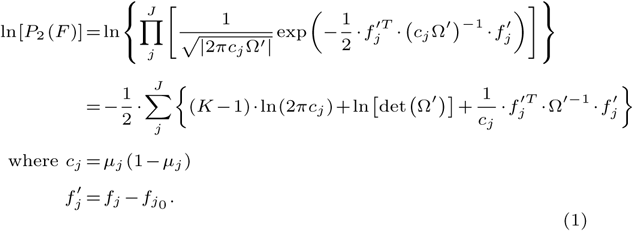

Following the structure analysis and component tree inference, a natural extension of this framework is to detect SNPs that deviate strongly from the globally estimated covariance structure. The idea of testing for deviations from a Gaussian distribution follows (Gnther and Coop, 2013), but differs in the use of an enforced tree-structure, an ML inference framework and fast optimization algorithms, thereby avoiding some of the computational challenges associated with Markov Chain Monte Carlo (MCMC). We also incorporate admixture into our model, thereby enabling the possibility to test for positive selection acting on the ancestral components of a panel, before more recent admixture occurred between the ancestors of the sampled individuals

### Selection model

The test for selection is based on a likelihood ratio test that identifies SNPs with allele frequency patterns that are poorly described by the genome-wide covariance pattern. A genome-wide covariance matrix is estimated from all SNPs jointly. Each SNP is then independently tested for deviations from this model, using a scalar factor introduced to certain elements of the covariance matrix. This scalar factor can be introduced in different ways depending on which selection hypotheses are tested. In our analyses, we chose to scale the covariance matrix such that one of its diagonal values is multiplied by a scalar, *α*, corresponding to strong differences in allele frequency in one of the ancestry components relative to the rest:

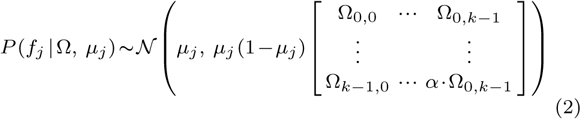

The value of *α* is then estimated using ML and a likelihood ratio is formed by testing the hypothesis of *α* = 1 against the alternative of *α* > 1. A high likelihood ratio indicates a larger deviation in allele frequency in a focal component than expected under the globally estimated null-model. Figure 1 shows an example. This test can also be implemented to test selection on ancestral non-terminal lineages by multiplying the corresponding values in the covariance matrix by a scaling factor.

**FIG. 1.**
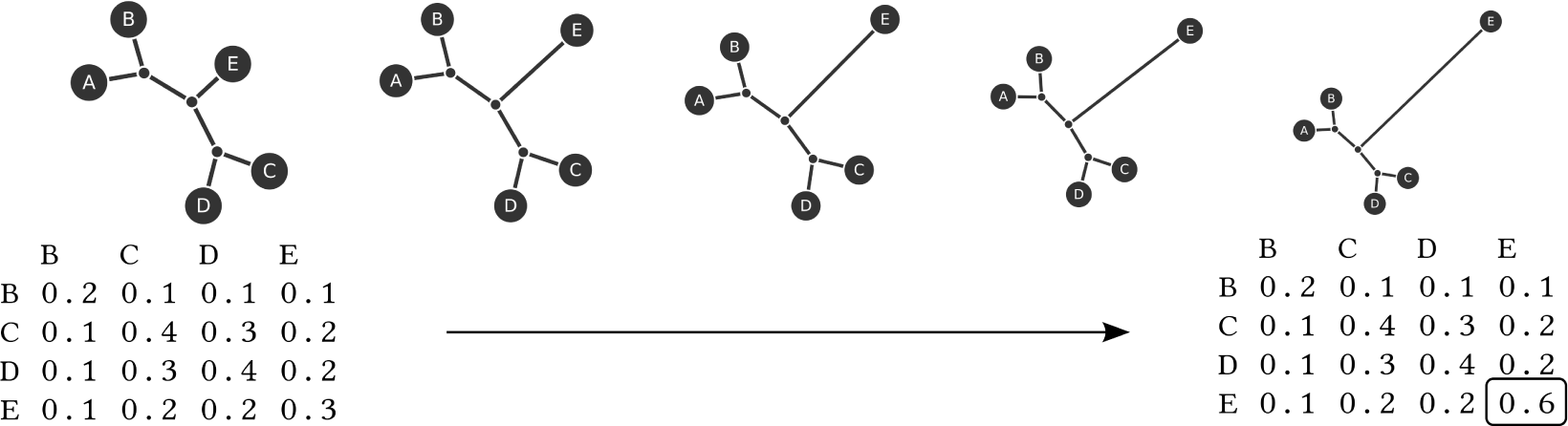
Selection hypotheses and their encodings as covariance matrices. In this example, the ancestry component E is assumed to be the potential target of selection. The entry E:E in the covariance matrix is therefore allowed to deviate from the globally estimated value.

Under the null-hypothesis, the likelihood ratio test statistic is expected to follow a 50:50 mixture between a 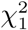-distribution and a point mass at zero (Self and Liang, 1987).

In summary, we estimate a scaling factor for one or more components of the covariance matrix in a multivariate normal model of allele frequency distribution among populations. For each candidate SNP, we then compare the estimated covariance matrix to that obtained genome-wide, using a likelihood ratio test.

### Optimization

For ML-based population structure inference, we use an optimization algorithm based on an Active Set method (Murty and Yu, 1988) to solve the sequential quadratic programming problem. This method was previously shown to have better computational performance than competing methods (Cheng *et al.*, 2017). For the ML-based ancestry covariance estimates, we use the Nelder-Mead simplex method (Nelder and Mead, 1965). It uses Cholesky decomposition (Cholesky, 1910) to determine the positive semi-definiteness of a matrix and to compute matrix inverses and determinants. For identifying the best local covariance structure during a selection scan, we use a simple Golden-section search algorithm (Kiefer, 1953) to find the solution for the single scalar multiplier associated with a specific selection hypothesis.

### Simulations

To evaluate the performance of the methods, we generate simulations using *msms* (Ewing and Hermisson, 2010) under specific demographic models and specific tree structures (Figure 2). We focus on multi-population demographics that are simulated in a tree-like fashion with positive selection events occurring in either all or some of the branches leading to present-day samples.

**FIG. 2.**
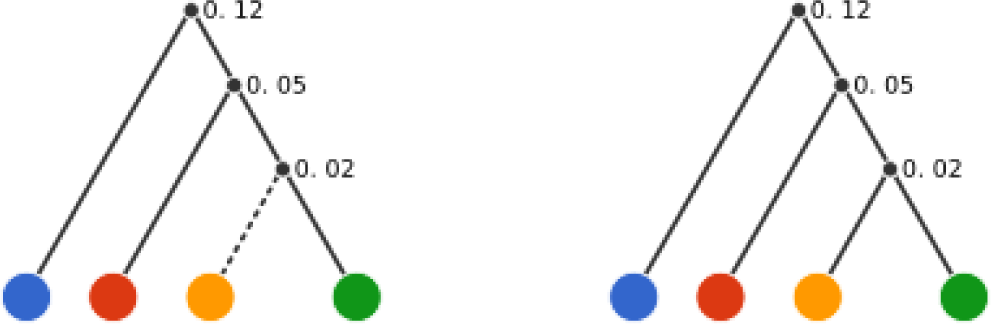
Trees used for simulations. We simulated selection only on the yellow branch (left) and also simulated neutral segments according to the same tree but with no selection on any branches (right).

Specifically, we simulate an effective population size *N*_*e*_ of 10,000 for all populations, and obtain 20 chromosomes for each population. We use a population-scaled mutation rate *θ* = 4*N*_*e*_*µ* of 100, a recombination rate *ρ* = 4*N*_*e*_*r* of 100, with an finite cut site model as implemented in MSMS. We simulate 4 populations with population splits at 0.02, 0.05, and 0.12 coalescent units in the past (in units of 4*N*_*e*_). This is illustrated in Figure 2.

We assessed the power of our selection test using simulations. We simulated 1,200 bp sequences where a single beneficial SNP located in the middle of the sequence is under direct positive selection. We also simulated 1,200 bp neutral sequences. We set the start of selection 0.02 coalescent units ago, and set the initial frequency of the beneficial allele at 0 (i.e. we simulate a *de novo* mutation). We assumed an additive model of fitness with scaled selection coefficient 2*N*_*e*_*s* ranging from 200 to 1000 for alleles in the homozygous state. In our model, the fitness of the heterozygote is 1+*s/*2 and the two homozygous fitnesses are 1 and and 1+*s*. We set the forward mutation rate, 4*N*_*e*_*µ*′, to 0.1 for the selected allele for mutations from the wild type to the selected type (the backwards mutation rate to the wild type is 0). In the simulations used for Figure 5, we combined 2 neutral sequences flanking a selected sequence. In the simulations used for Figure 4, we placed 10 neutral sequences flanking a selected sequence on either side.

A population can then be purely formed by one ancestry component, or as a mixture of several components. We generate un-admixed or admixed genotypic data by simulating the ancestry proportions, *Q*, for each individual (Figure 3). For un-admixed samples, we simply assign them to have 100% ancestry from one population. For admixed samples, we simulate equal admixture (in expectation) using the Dirichlet distribution, Dir(*α*), where *α* =(1.0,*…*,1.0). Figure 3 illustrates these two admixture setups. In the mixed case, each population is (in expectation) an equal mixture of three of the ancestry components, i.e. each population lacks one of the four components.

**FIG. 3.**
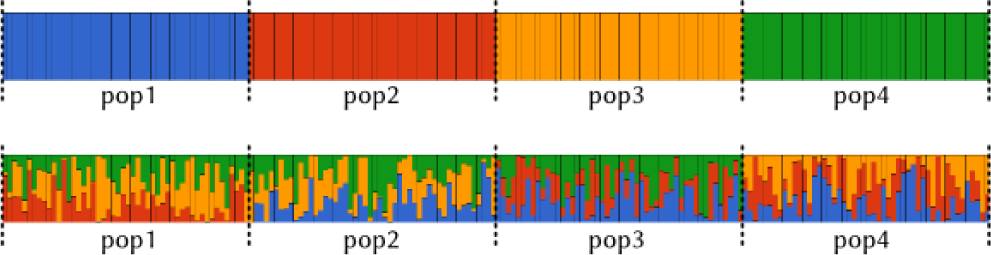
Simulated admixture proportions for either un-admixed individuals (top) or equal admixture from three out of four ancestry components (bottom). In both scenarios, we simulated 4 populations of 20 individuals per population.

We then sampled genotype observations under the assumption of independence, i.e.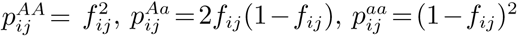, where *f*_*ij*_ =*Σ*_*k*_ *Q*_*ik*_*F*_*kj*_ is the allele frequency in locus *j* for individual *i, Q*_*ik*_ is the ancestry proportion of component *k* in individual *i, F*_*kj*_ is the allele frequency in ancestry component *k* in locus *j*, and *p*_*AA*_, *p*_*Aa*_ and *p*_*aa*_ are the probabilities of observing major-major, major-minor, or minor-minor genotypes for the locus, respectively. *F* has dimensionality *K × J* and *Q* has dimensionality *I × K*, where *K* is the number of ancestry components, *I* is the number of samples, and *J* is the number of SNPs.

## Results

### Simulations

We first evaluated the efficacy of the method for fine-mapping the true selected allele. We simulated 1000 replicates of 4 admixed populations and scanned the simulated genomes using our likelihood ratio test. As a measure of accuracy, we used the distance between the SNP with the highest likelihood ratio and the SNP under selection in the simulations. In the majority of simulations, the distance between the true and the inferred SNP is small, i.e. *<* 10% of sequence length, suggesting a generally high accuracy for fine-mapping.

We then measured the excess of false positive results under the null hypothesis of no selection (Figure 4). To do so, we first generated simulations under the null scenario (no selection, Figure 4-top) and a scenario of positive selection affecting a particular ancestry branch (Figure 4-bottom). We also simulated 2 types of samples: un-admixed (left) and admixed (right). We computed per-SNP likelihood ratios along the simulated genomes using the correct selection model and converted them to p-values. We then compared these p-values in a quantile-quantile (QQ) plots against a 50:50 mixture of values equal to 1 and random values sampled from a uniform distribution between 0 and 1. This mixture corresponds to the expected distribution of P-values of our statistic under the null model. In simulations with selection, deviations from the neutral expectation are visible (Figure 4-bottom), while no excess of false positives (elevated Type I errors) are present when populations are simulated without selection (Figure 4-top).

**FIG. 4.**
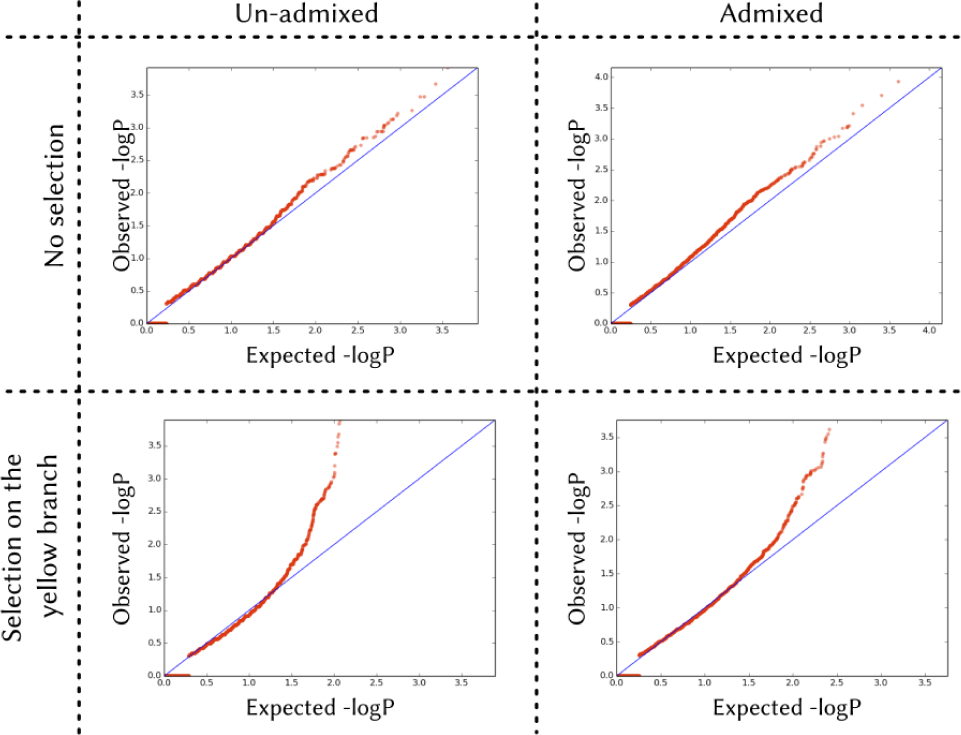
Test for excess of false positive. We simulate 4 different scenarios: no selection (top) and selection on a single branch (bottom), assuming the input populations are un-admixed (left) and admixed (right). Statistically significant outliers are detected only when the data were simulated under the scenarios with selection.

We then compared our method to three summary statistics: *F*_*ST*_ (Wright *et al.*, 1949; Weir *et al.*, 2005), PBS (Yi *et al.*, 2010), and FLK (Bonhomme *et al.*, 2010) (Figure 5). As before, we generated simulations under 2 types of sample admixture, and simulated positive selection on the yellow component only. We tested for selection using 9 methods: Ohana, 2 *F*_*ST*_-based tests statistics, 2 PBS-based tests, and 4 FLK-based tests. The Ohana method was run so as to test specifically for selection on the yellow-branch. In one of the *F*_*ST*_-based tests, we calculated pairwise *F*_*ST*_ between 2 populations: one containing the yellow component and one not containing it. In the second *F*_*ST*_-based test, we calculated *F*_*ST*_ among all 4 populations. In the 2 PBS-based tests, we tested for selection focusing on either the yellow or the green component as the “target” branch. In the 4 FLK-based tests, we specified each of the 4 populations as the outgroup in turn.

**FIG. 5.**
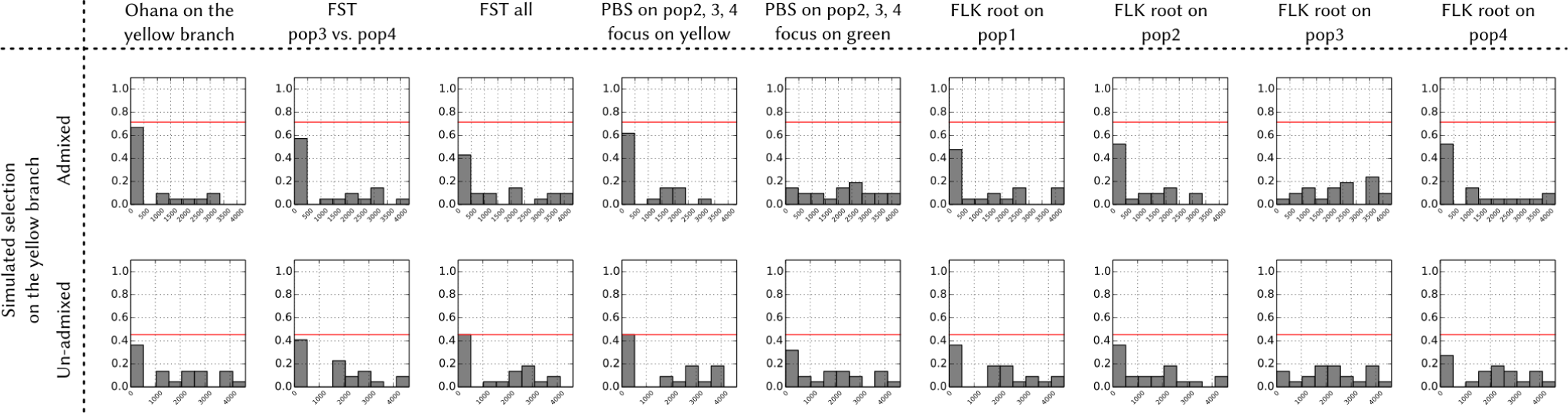
Comparison of performance of selection tests. We simulated individuals that were either admixed (row 1) or un-admixed (row 2), as described in Figure 3. We then simulated selection on a single branch (yellow). We simulated the SNP in the midpoint location to be under direct selection, with a selection strength of 2*N*_*e*_*s* = 600 for the homozygote and 2*N*_*e*_*s* = 300 for the heterozygote. We then scanned for selection signals using Ohana (first column). We calculated the FST statistics in 2 ways: FST between two population (one with the yellow component one without), and FST among all four populations. We also calculated the PBS statistics in 2 ways: yellow-specific or green-specific. Finally, we calculated the FLK statistics in 4 ways by specifying each of the 4 populations as the outgroup. In admixed simulations, Ohana outperforms the rest of the tests by achieving a higher proportion of simulations in which the simulated SNP under selection is < 120*bp* (10% of total sequence length) from the SNP with the highest test score.

The performance of the methods was measured by the percentage of runs in which the simulated and detected SNP are within 10% of the total sequence length. In admixed samples, our method achieves the best outcome when the proper model is specified (Figure 5 row 1). When analyzing purely un-admixed samples, our method is on-par with the best method among all *F*_*ST*_, PBS, and FLK tests (Figure 5 row 2).

We then compared all methods in Figure 5 using 2 measures: the percentage of times that the selection method identifies the simulated causal SNP as the top SNP (Table 1) and the mean distance in bp between the simulated causal SNP and the top SNP identified by the selection method (Table 1 and 2).

**Table 1.**
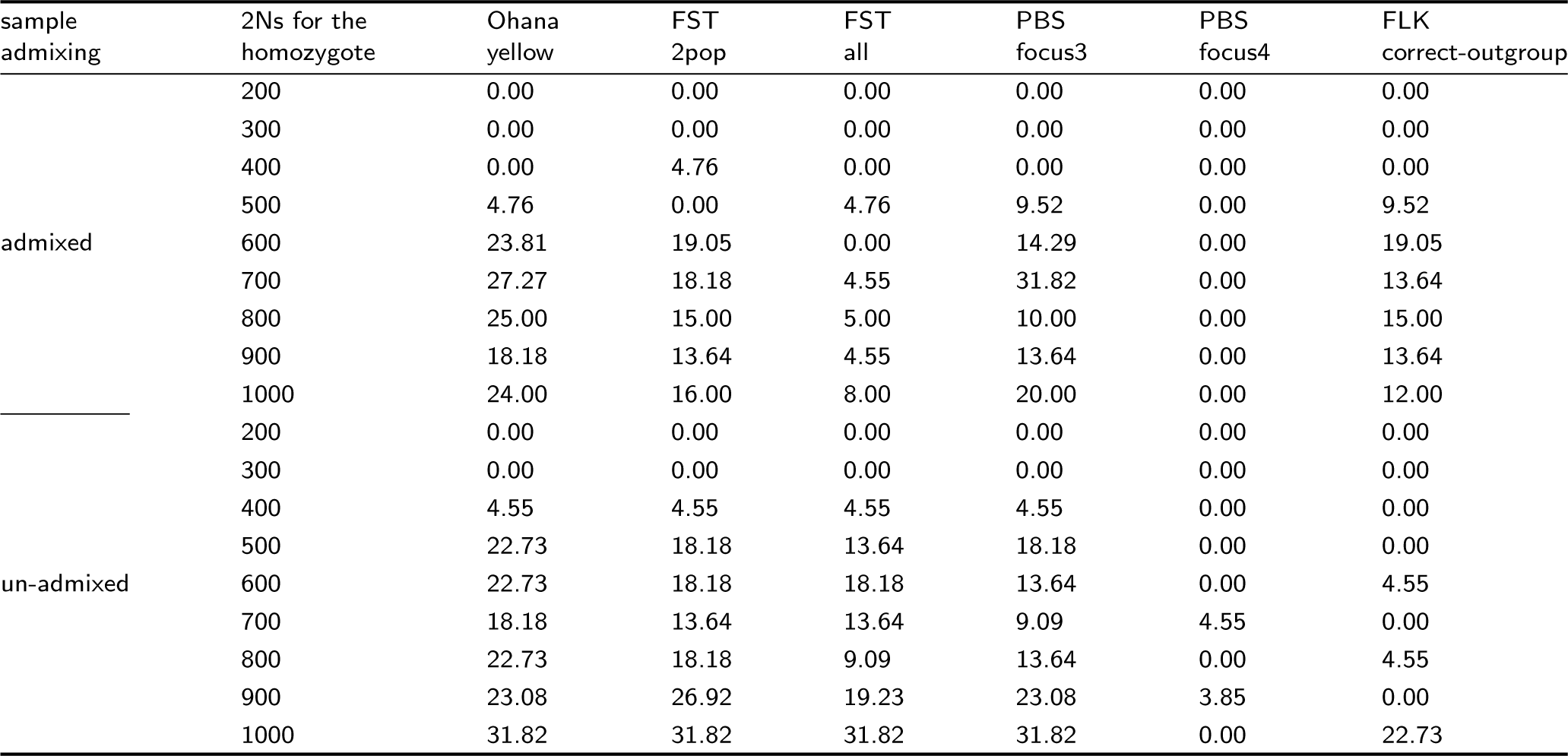
We compared the new method with the *F*_*ST*_, PBS, and FLK statistics. The data simulation and selection detection are as described in Figure 5. We quantify selection strength using the percentage of simulations among a total of 500 where the selection method accurately identifies the simulated causal SNP as the top SNP.

**Table 2.**
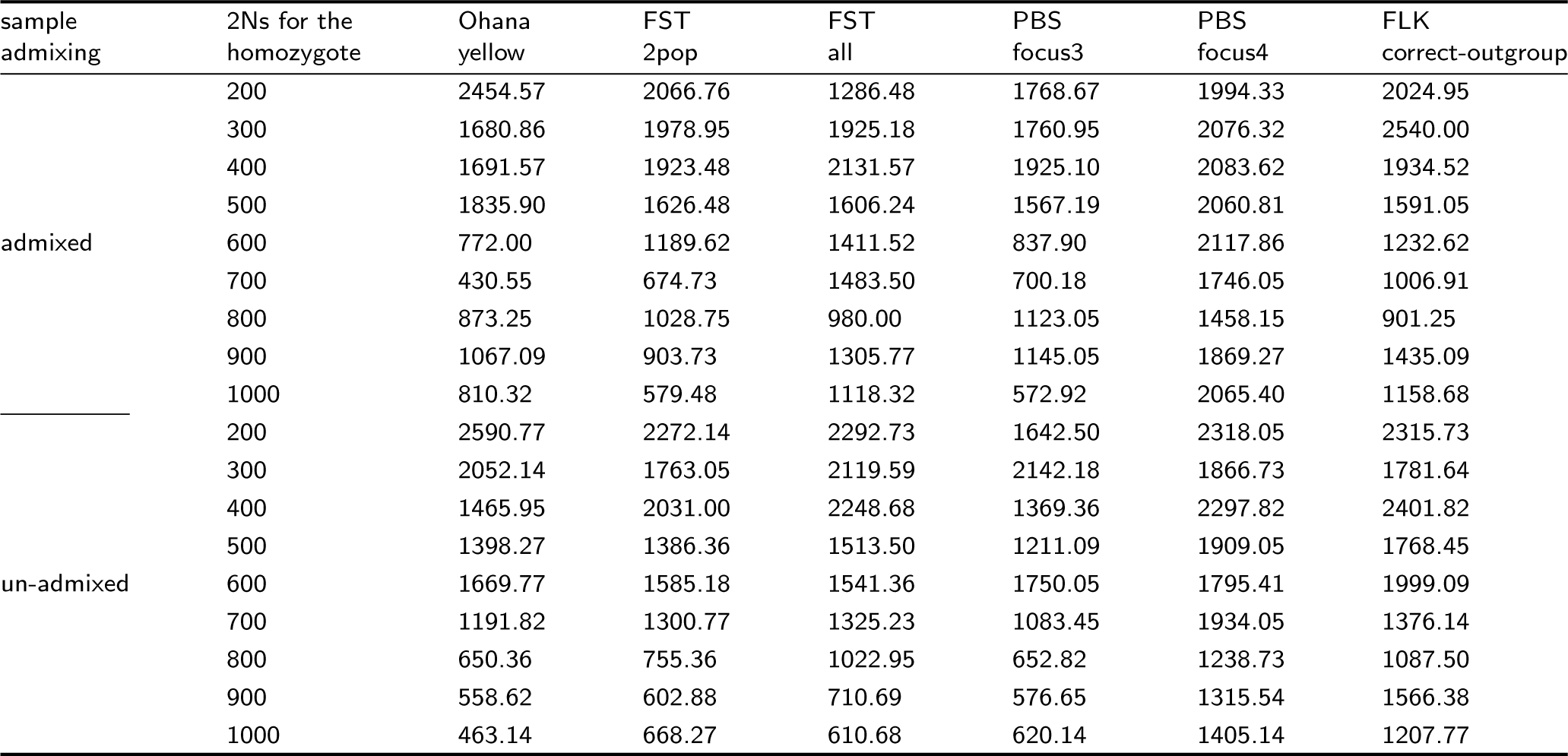
Mean distance between the top-scoring SNP and the simulated beneficial SNP. We compare the new method with the *F*_*ST*_, PBS and FLK statistics. The data simulation and selection detection methods are as described in Figure 5.

For all methods, scenarios with admixed samples lead to weaker performance than in their un-admixed counterparts. In un-admixed cases, our method performs equally or better than summary statistics. In admixed cases, our method reaches higher accuracy than other methods. For example, in the admixed yellow-branch simulation when 2*Ns* = 600, Ohana identifies the selected allele 21.57% of the time, while this value is 14.95% for pairwise *F*_*ST*_, 8.09% for global *F*_*ST*_, 18.63% for PBS, and 7.11% for FLK.

### Analysis of real data

We identified regions in the genome that are likely to have been under the influence of positive selection using a merged dataset containing several population panels from phase 3 of the 1000 Genomes Project (1000 Genomes Project Consortium, 2015). We randomly selected 64 genomes from each of 4 populations from the 1000 Genomes project: the British from Great Britain (GBR), the Han Chinese from Beijing (CHB), the Yoruba Africans (YRI) and the admixed Mexican-Americans from Los Angeles (MXL) (the number 64 was chosen because it was the size of the smallest panel). We only included variable sites with no missing data and a minimum allele frequency of 0.05 across the entire merged panel. In total, we analyzed 5,601,710 variable sites across the autosomal genome. We inferred genome-wide allele frequencies and covariances for the latent ancestry components as described in the Methods section, using *K* = 4. To scan for covariance outliers, we performed four hypothesis-driven scans, in which we specifically searched for selection separately in each of the four inferred ancestry components in our dataset (Table 3).

**Table 3.** Top 10 most differentiated SNPs from each of the ancestry-specific scans. LLRS = log-likelihood ratio score for positive selection.

| chr | pos | rsid | LLRS | target ancestry | nearest gene |
| --- | --- | --- | --- | --- | --- |
| 5 | 33951693 | rs16891982 | 22.085902 | European | SLC45A2 |
| 15 | 48426484 | rs1426654 | 19.707464 | European | SLC24A5 |
| 15 | 28356859 | rs1129038 | 19.290553 | European | HERC2 |
| 15 | 28495956 | rs12912427 | 18.270213 | European | HERC2 |
| 9 | 16792200 | rs10962596 | 15.819739 | European | BNC2 |
| 1 | 1385211 | rs1312568 | 15.066101 | European | ATAD3C |
| 2 | 136407479 | rs1446585 | 14.957582 | European | R3HDM1 |
| 2 | 136616754 | rs182549 | 14.629386 | European | MCM6 |
| 1 | 204784969 | rs3940119 | 14.393216 | European | NFASC |
| 4 | 38798648 | rs5743618 | 14.38681 | European | TLR1 |
| 16 | 48258198 | rs17822931 | 23.271759 | East Asian | ABCC11 |
| 16 | 48375777 | rs6500380 | 22.474103 | East Asian | LONP2 |
| 1 | 234635790 | rs2175591 | 20.95541 | East Asian | TARBP1 |
| 4 | 100142780 | rs75721934 | 20.453247 | East Asian | LOC100507053 |
| 11 | 61579427 | rs72643557 | 20.114033 | East Asian | FADS1 |
| 11 | 120154631 | rs12224052 | 19.696284 | East Asian | POU2F3 |
| 21 | 43974948 | rs228088 | 19.518001 | East Asian | SLC37A1 |
| 11 | 133043841 | rs79802711 | 19.157192 | East Asian | OPCML |
| 5 | 128016573 | rs79478220 | 18.476104 | East Asian | FBN2 |
| 19 | 51441759 | rs11084040 | 18.158963 | East Asian | KLK5 |
| 14 | 46745012 | rs140736443 | 32.730697 | Native American | LINC00871 |
| 9 | 82968379 | rs6559543 | 27.584847 | Native American | LINC01507 |
| 16 | 80619307 | rs2316155 | 27.399123 | Native American | LINC01227 |
| 14 | 21647765 | rs77549780 | 27.355769 | Native American | LINC00641 |
| 12 | 14189549 | rs12425115 | 25.867367 | Native American | GRIN2B |
| 10 | 8150713 | rs10508343 | 25.609772 | Native American | GATA3 |
| 15 | 34936250 | rs16959274 | 25.424824 | Native American | GOLGA8B |
| 8 | 4490837 | rs71523639 | 24.59957 | Native American | CSMD1 |
| 1 | 14301862 | rs72640512 | 24.455822 | Native American | PRDM2 |
| 12 | 29817716 | rs12580697 | 23.967094 | Native American | TMTC1 |
| 14 | 57318110 | rs75607199 | 23.959875 | Native American | OTX2-AS1 |
| 8 | 145639681 | rs1871534 | 11.906794 | Yoruba / Ancestral Non-African | SLC39A4 |
| 5 | 178626609 | rs6869589 | 11.541667 | Yoruba / Ancestral Non-African | ADAMTS2 |
| 15 | 29427400 | rs10152250 | 11.48232 | Yoruba / Ancestral Non-African | FAM189A1 |
| 1 | 1106112 | rs6670693 | 11.447873 | Yoruba / Ancestral Non-African | TTLL10 |
| 4 | 3666494 | rs58827274 | 11.341367 | Yoruba / Ancestral Non-African | LOC100133461 |
| 17 | 2631985 | rs4790359 | 11.118134 | Yoruba / Ancestral Non-African | PAFAH1B1 |
| 9 | 136769888 | rs2789823 | 11.031687 | Yoruba / Ancestral Non-African | VAV2 |
| 6 | 169656029 | rs6930377 | 10.824098 | Yoruba / Ancestral Non-African | THBS2 |
| 17 | 29350769 | rs8073072 | 10.794224 | Yoruba / Ancestral Non-African | RNF135 |
| 5 | 173642871 | rs10067518 | 10.787147 | Yoruba / Ancestral Non-African | HMP19 |

After running these scans, we queried the CADD server (Rentzsch *et al.*, 2018) to obtain functional, conservation and regulatory annotations for the top candidate SNPs, including SIFT (Sim *et al.*, 2012), PolyPhen (Adzhubei *et al.*, 2013), GERP (Davydov *et al.*, 2010), PhastCons (Siepel *et al.*, 2005), PhyloP (Pollard *et al.*, 2010) and Segway (Hoffman *et al.*, 2012) annotations, so as to find the changes most likely to be disruptive. We discuss some of these below. We also queried the GTEx cis-eQTL database (Lonsdale *et al.*, 2013), the UK Biobank GeneATLAS (Canela-Xandri *et al.*, 2018), and the GWAS catalog (MacArthur *et al.*, 2017), to look for trait-associated SNPs. We particularly focus on SNPs that have both high log-likelihood ratios in favor of positive selection (*LLRS >* 15) and high CADD scores in favor of functional disruption (*>* 10).

Below, we describe some of the top SNPs with high LLRS and their surrounding regions, for those cases in which available genic, expression or regulatory information can provide us some clue as to the possible organismal function that may have been affected by the selective event. We particularly focus on the Native American ancestry scan (Table S4, Figure 6), as few selection scans have been performed in this population, but also briefly summarize the results from the other scans.

**FIG. 6.**
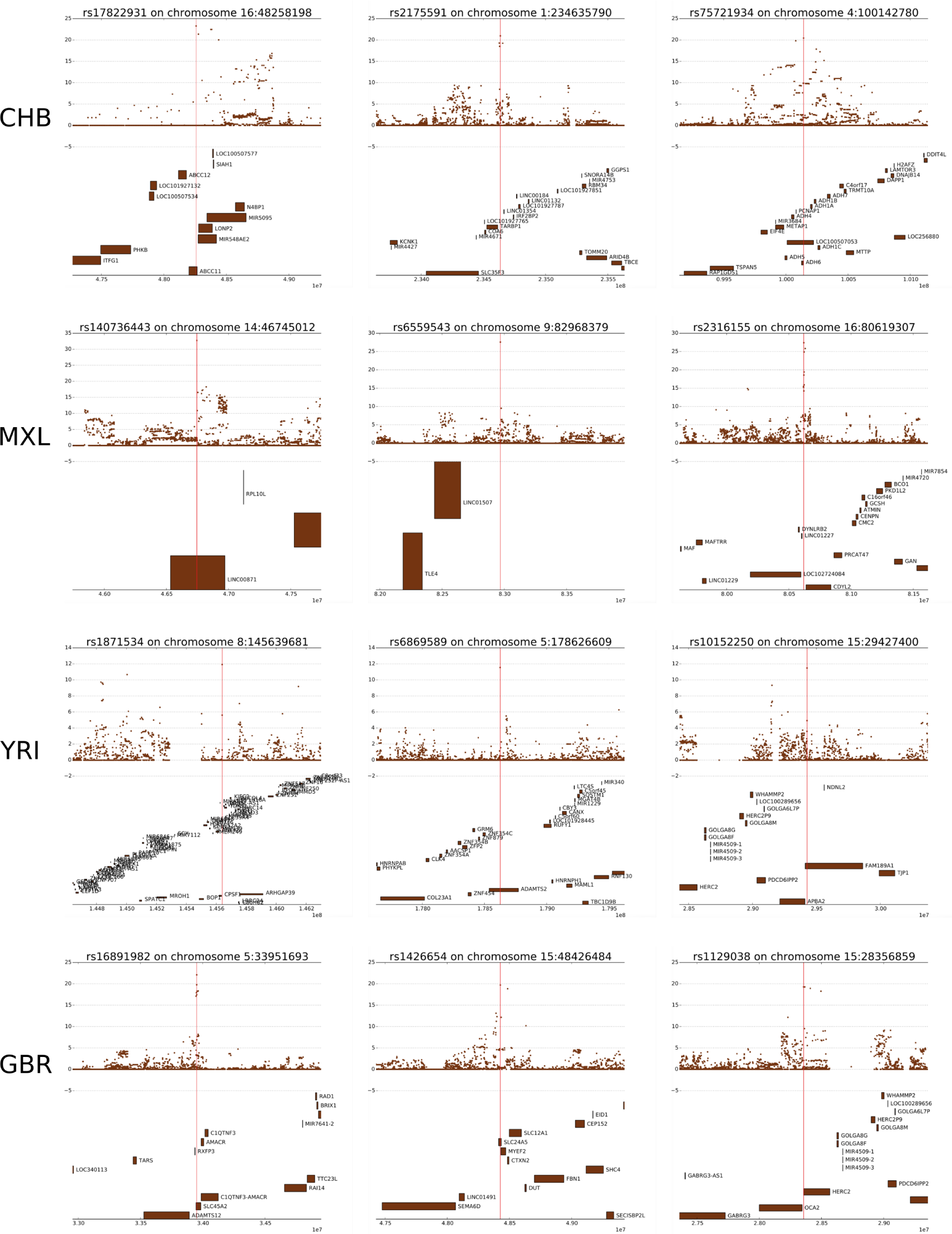
Top 5 annotated peaks in each of the ancestry-specific selection studies. MXL-specific = scan for selection in Native American ancestry of MXL. GBR-specific = scan for selection in European ancestry of GBR. CHB-specific = scan for selection in East Asian ancestry of CHB. YRI-specific = scan for selection in Yoruba African ancestry or ancestral non-African ancestry. We analyzed 5,601,710 variable sites across the autosomal genomes. We inferred genome-wide allele frequencies and covariances as described in the Methods section. We applied a likelihood model for each SNP by rescaling all variances and covariances by a scalar multiplier *α*. Descriptions of each candidate region are in Table 3.

### European ancestry scan

Results for the top 30 loci in the European ancestry scan are presented in Table S1. Most loci have been previously shown to be under selection in Europeans populations, including SLC45A2, SLC24A5, BNC2, the OCA2/HERC2 region, the LCT/MCM6 region and the TLR region (Mathieson *et al.*, 2015;Bersaglieri *et al.*, 2004;Voight *et al.*, 2006;Vernot *et al.*, 2014;Barreiro *et al.*, 2009). We notice that, in several cases, the presumed causal SNP previously identified in the literature coincides with the SNP with the strongest selection signal. This is the case, for example, for rs1426654 (SLC24A5) (Kimura *et al.*, 2009;Lamason *et al.*, 2005) and for rs16891982 (SLC45A2) (Branicki *et al.*, 2008). This suggest that the top SNPs for other loci, for which the causal SNPs are not yet known, may be good candidates for further tests of functional effects.

### East Asian ancestry scan

We also performed a scan where we sought to recover SNPs that were candidates for selection in the ancestry component that is prevalent among our East Asian samples. Results for the top 30 loci in this scan are in Table S2. Here, we also recover several candidate regions that have been previously reported in East Asian selection scans, including ABCC11, POU2F3, ADH1B, FADS1 and TARBP1 (Vernot *et al.*, 2014;Liu *et al.*, 2018;Ohashi *et al.*, 2011;Peng *et al.*, 2010;Refoyo-Martinez *et al.*, 2018). Here, as in the previous scan, the top-scoring SNPs also tend to have the strongest phenotypic associations. For example, the highest scoring SNP (rs17822931) is the well-known missense variant in ABCC11, which is involved in sweat and earwax production (Yoshiura *et al.*, 2006).

### Yoruba / ancestral non-African ancestry scan

Because our algorithm relies on an unrooted ancestry tree, we cannot distinguish between SNPs under positive selection in the terminal branch leading to the Yoruba / Sub-Saharan Africans and the ancestral non-African branch (Table S3). Nevertheless, more careful study of the allele frequencies of these SNPs in other populations may serve to distinguish among these scenarios in the future. As in the the other ancestry scans, we also retrieve several genes that have been previously reported in positive selection studies. For example, the highest-scoring SNP is a missense variant in SLC39A4 (rs1871534) that has been reported to be under selection in Sub-Saharan Africa and to be causal for zinc deficiency (Engelken *et al.*, 2014).

### Native American ancestry scan

The Native American ancestry scan yielded several novel candidates for positive selection (Table S4). As this ancestry has been less studied than the other aforementioned populations in the selection scan literature, we decided to extensively describe the top 30 candidates in the Supplementary Notes. We also highlight some of the more interesting regions here.

The top SNP (rs140736443) is located in an intron of LINC00871. This SNP does not have a high CADD score (= 1.125), but is very close to a SNP (rs10133371) with a very high LLRS (= 16.54) and CADD score (= 15.99). This SNP is also intronic but is highly conserved in primates (PhastCons = 0.972) and is located in a GERP conserved element (P = 1.92e-21). LINC00871 is a long non-coding RNA gene that has been associated with number of children born (Barban *et al.*, 2016), although the specific trait-associated SNP in that study does not have a high LLRS. This gene also contains a suggestive association to longevity in females (Zeng *et al.*, 2018), although this study was under-powered to retrieve genome-wide significant associations.

The third top SNP (rs2316155) has a low CADD score (= 0.633) but is located near two SNPs with high LLRS (rs1466182, rs1466183) that overlap a regulatory region (ENSR00000088366) and have high CADD scores (= 16.8 and 19.5, respectively). Both of these SNPs have high PhastCons conservation scores across primates, mammals and vertebrates, and both overlap a GERP conserved element.

The sixth top SNP (rs10508343) has a low CADD score but lies very close to another SNP (rs17143255) with a high LLRS and a very high CADD score (= 14.16). The latter is an intergenic SNP overlapping a GERP conserved element between LINC00708 and GATA3, which has been shown to lead to abnormal hair shape and growth in mice when mutated (Kaufman *et al.*, 2003). Interestingly, SNPs overlapping LINC00708 have been recently associated with hair shape in a GWAS of admixed Latin Americans (Adhikari et al. 2016). There is also a high-LLRS SNP in this region that is significantly associated with the response to treatment for acute lymphoblastic leukemia (rs10508343) (Yang *et al.*, 2009).

The seventh top SNP (rs16959274) is a GTEx eQTL for GOLGA8A for tibial artery and skeletal muscle, and for GOLGA8B in pancreas. These two genes are members of the same gene family, and code for an auto-antigen localized in the surface of the Golgi complex (Eystathioy *et al.*, 2000).

The tenth top SNP (rs12580697) is a GTEx eQTL for TMTC1 in whole blood and has a moderately high CADD score (= 8.676). TMTC1 codes for an endoplasmic reticulum transmembrane protein that is involved in calcium homeostasis (Sunryd *et al.*, 2014).

The eleventh top SNP (rs75607199) has a low CADD score but lies near three other SNPs (rs41325445, rs4901738 and rs59250732) with almost equally high LLRS and high CADD scores (= 13.49, 19.7 and 12.67, respectively). All of these SNPs are intronic and overlap OTX2-AS1, a long non-coding RNA gene. The SNP with the highest CADD score (rs4901738) is located in a GERP conserved element and has high PhastCons conservation scores across primates and mammals (*>* 0.98). They all lie upstream of OTX2, coding for a developmental transcription factor implicated in microphtalmia (Ragge *et al.*, 2005), retinal dystrophy (Vincent *et al.*, 2014) and pituitary hormone deficiency (Diaczok *et al.*, 2008). In mice, this gene has been found to be involved in the embryonic development of the brain (Boncinelli *et al.*, 1993), photoreceptor development (Nishida *et al.*, 2003) and susceptibility to stress (Peña *et al.*, 2017).

The fourteenth top SNP (rs78441257) has a fairly high CADD score (= 12.72) and lies in a GERP conserved element of the 3’ UTR of LRAT. This gene is implicated in retinal dystrophy (Thompson *et al.*, 2001) and retinitis pigmentos (Sénéchal *et al.*, 2006).

The fifteenth top SNP (rs1919550) is a GTEx eQTL for FBXO40 in whole blood, but does not have a high CADD score. However, it lies near a SNP (rs9813391) with a high LLRS that leads to a nonsynonymous change (R145Q) in ARGFX-a homeobox gene- and another SNP (rs4676737) with both a high LLRS and high CADD score (= 14.07) overlapping a repressor region in an intron of FBXO40. The latter SNP is a GTEx eQTL for IQCB1 in fibroblasts, muscular esophagus and thyroid. IQCB1 is associated with Senor-Loken syndrome (Otto *et al.*, 2005), a ciliopathic eye disorder.

The twenty-second top SNP (rs4946567) is an eQTL of TBC1D32 in cerebellar brain. This SNP has a high CADD score (= 11.02) and is conserved across vertebrates (vertebrate PhyloP = 0.916, vertebrate PhastCons = 0.747). Interestingly, the region in which it is located also harbors signature of selection in Yucatan miniature pigs (Kim *et al.*, 2015;Kwon *et al.*, 2018). TBC1D32 plays a role in cilia assembly (Ko *et al.*, 2010) and may be involved in ciliopathic congenital abnormalities, including midline cleft, microcephaly, and microphthalmia (Adly *et al.*, 2014).

The twenty-third and twenty-fourth top SNPs (rs5758430, rs4822061) are close to each other and lie in a large region with several high-LLRS SNPs. They are both linked GTEx eQTLs to several genes in a variety of different tissues. They are also both significantly associated with several traits related to body fat, food intake and white blood cells in the UK Biobank GeneATLAS (P *<* 10^-8^, see Supplementary Notes). Although these SNPs do not have particularly high CADD scores, there are several neighboring linked high-LLRS, high-CADD SNPs with significant associations to the same traits, including splice site and missense mutations (Supplementary Notes). We also find two significantly-associated SNPs in the GWAS catalog in this region (P *<* 10^-8^): rs4822024 is associated with Vitiligo (Jin *et al.*, 2012) and rs13054099 is associated with neuroticism (Nagel *et al.*, 2018).

## Discussion

We describe a new modeling framework that can detect signals of positive selection on ancestry components, using allele frequency patterns across admixed populations. It models admixture explicitly and works with an arbitrary number of populations with or without admixed ancestries. It also does not rely on labeling of samples into particular populations, and allows for testing of different positive selection models reflecting different historical adaptive hypotheses.

The run-time complexity of our method is linear in the number of markers, but we still recommend a high-performance cluster to be used in a typical genomic analysis. With parallelization, a selection scan takes *<* 10 minutes to analyze a 6 Mbp genome for *<* 10 ancestry components using 100 cores. An example of how to perform this parallelization can be found on the project’s wiki page on GitHub.

Our method works by testing for selection in specific components of the ancestry covariance matrix. We also explored what would occur if we used a likelihood model in which the ancestry covariance matrix was multiplied by a scalar, so as to find “global” candidates for selection rather than testing for selection in particular ancestries. We found however, that this was not an optimal way to detect candidates for selection, as it is biased towards finding many variants in highly drifted populations, likely because the excess variance in the Wright-Fisher process is not well modelled by the multivariate Gaussian assumption, especially at the boundaries of fixation and extinction.

When specifically testing for candidates for selection in Europeans, East Asians and Sub-Saharan Africans we identified several well-known candidates under positive selection, including OCA2, SLC24A5, SLC45A2, ABCC1 and SLC39A4. Many of our top scoring SNPs were also previously known to be causal for particular traits, as in the case of rs17822931 in ABCC11 in East Asians, rs16891982 in SLC45A2 in Europeans, rs1426654 in SLC24A5 in Europeans and rs1871534 in SLC39A4 in Sub-Saharan Africans.

Our scan for positive selection in the Native American ancestry component of Latin Americans yielded several novel candidates for adaptation in the human past. We found signatures of selection near genes involved in fertility (LINC00871), hair shape and growth (LINC00708), immunity (GOLGA8A / GOLGA8B and IRAK4), vision (OTX2 and LRAT), the nervous system (MDGA2) and various ciliopathies (IQCB1 and TBC1D32). Several of the highest-scoring SNPs in the candidate regions are known to be cis-eQTLs to their nearby genes, as is the case for rs12580697 / TMTC1 (involved in calcium homestasis) and rs4676737 / IQCB1 (involved in ciliopathies). We also found individual SNPs with high likelihood ratio scores in favor of selection that are associated with a variety of phenotypes, including rs12426688 (fat percentage), rs10508343 (response to leukemia treatment), rs34670506 (insomnia), and the cluster of high-scoring SNPs that include rs5758430 and rs4822061, among other SNPs. This particular cluster is especially interesting, as the SNPs in the region are associated with a variety of traits related to body fat distribution, food intake and white blood cells, suggesting a possible underlying phenotype related to these traits that may have driven an adaptive event.

We provide a list of functional annotations for all the SNPs with high LLRS (*>* 15) within a 2Mb region surrounding each of the top genome-wide SNPs, including CADD, conservation, regulatory and protein deleteriousness scores, which we hope will guide future functional validation studies in these regions of the genome (Table S5).

In conclusion, Ohana provides a fast and flexible selection-detection and hypothesis-testing framework. It is easy to use and has in-built visualization functionalities to explore patterns on a genome-wide and locus-specific scale. We believe it will be a useful tool for biologists aiming to study positive selection and understanding the genomic basis of adaptation, particularly in cases where demographic histories are complex or not well characterized.

## Acknowledgments

The authors gratefully acknowledge Thomas Mailund, Mikkel Schierup, Christian Storm Pedersen, and the GenomeDK staff for their support during the course of this research. FR thanks the Villum Foundation for their support.

