## supplement for "Ohana: detecting selection in multiple populations by modelling ancestral admixture components"

February 11, 2019

### Candidates for selection in Native American ancestry

- The top SNP (rs140736443) is located in an intron of LINC00871. This SNP does not have a high CADD score (= 1.125), but is very close to a SNP (rs10133371) with a very high LLRS (= 16.54) and CADD score (= 15.99). This SNP is also intronic but is highly conserved in primates (PhastCons = 0.972) and is located in a GERP conserved element (P = 1.92e-21). LINC00871 is a long non-coding RNA gene that has been associated with number of children born (Barban *et al.*, 2016), although the specific trait-associated SNP in that study does not have a high LLRS. This gene also contains a suggestive association to longevity in females (Zeng *et al.*, 2018), although this study was under-powered to retrieve genome-wide significant associations.

- The second top SNP (rs6559543) is near the genes LINC01507 and TLE4. It has a low CADD score (= 0.015) and no other nearby SNPs with high LLRS.

- The third top SNP (rs2316155) has a low CADD score (= 0.633) but is located near two SNPs with high LLRS (rs1466182, rs1466183) that overlap a regulatory region (ENSR00000088366) and have high CADD scores (= 16.8 and 19.5, respectively). Both of these SNPs have high PhastCons conservation scores across primates, mammals and vertebrates, and both overlap a GERP conserved element.

- The fourth top SNP (rs77549780) is an intergenic SNP downstream of CTD-2552B11.3, but does not have a high CADD score (= 1.348). Some of the nearby SNPs in the region have high LLRS scores with moderately high CADD scores, but CTD-2552B11.3 is classified as a processed pseudogene with unknown function.

- The fifth top SNP (rs12425115) is very close to another SNP with high LLRS (rs12426688). They both have low CADD scores, but also a suggestive association for height and leg fat percentage in GeneATLAS ($P < 10^{-5}$).

- The sixth top SNP (rs10508343) has a low CADD score but lies very close to another SNP (rs17143255) with a high LLRS and a very high CADD score (= 14.16). The latter is an intergenic SNP overlapping a GERP conserved element between LINC00708 and GATA3, which has been shown to lead to abnormal hair shape and growth in mice when mutated (Kaufman *et al.*, 2003). Interestingly, SNPs overlapping LINC00708 have been recently associated with hair shape in a GWAS of admixed Latin Americans (Adhikari *et al.*, 2016). There is also a high-LLRS SNP in this region that is significantly associated with the response to treatment for acute lymphoblastic leukemia (rs10508343) (Yang *et al.*, 2009).

- The seventh top SNP (rs16959274) is a GTEx eQTL for GOLGA8A for tibial artery and skeletal muscle, and for GOLGA8B in pancreas. These two genes are members of the same gene family, and code for an auto-antigen localized in the surface of the Golgi complex (Eystathioy *et al.*, 2000).

- The eigth top SNP (rs71523639) is an intronic SNP in CSMD1 and has a low CADD score (= 0.035).

- The ninth top SNP (rs72640512) is an intergenic SNP lying downstream of PRDM2, and has a low CADD score (= 0.791).
- The tenth top SNP (rs12580697) is a GTEx eQTL for TMTC1 in whole blood and has a moderately high CADD score (= 8.676). TMTC1 codes for an endoplasmic reticulum transmembrane protein that is involved in calcium homeostasis (Sunryd *et al.*, 2014).
- The eleventh top SNP (rs75607199) has a low CADD score but lies near three other SNPs (rs41325445, rs4901738 and rs59250732) with almost equally high LLRS and high CADD scores (= 13.49, 19.7 and 12.67, respectively). All of these SNPs are intronic and overlap OTX2-AS1, a long non-coding RNA gene. The SNP with the highest CADD score (rs4901738) is located in a GERP conserved element and has high PhastCons conservation scores across primates and mammals (> 0.98). They all lie upstream of OTX2, coding for a developmental transcription factor implicated in microphtalmia (Ragge *et al.*, 2005), retinal dystrophy (Vincent *et al.*, 2014) and pituitary hormone deficiency (Diaczok *et al.*, 2008). In mice, this gene has been found to be involved in the embryonic development of the brain (Boncinelli *et al.*, 1993), photoreceptor development (Nishida *et al.*, 2003) and susceptibility to stress (Peña *et al.*, 2017).
- The twelfth top SNP (rs112850342) is in an intergenic region, lying upstream of EPCAM - associated with congenital enteropathy (Schnell *et al.*, 2013) and colorectal cancer (Ligtenberg *et al.*, 2009) - and CALM2 - associated with cardiac arrhythmia (Makita *et al.*, 2014).
- The thirteenth top SNP (rs13202771) lies in an intron of TIAM2.
- The fourteenth top SNP (rs78441257) has a fairly high CADD score (= 12.72) and lies in a GERP conserved element of the 3' UTR of LRAT. This gene is implicated in retinal dystrophy (Thompson *et al.*, 2001) and retinitis pigmentosa (SÃľnÃľchal *et al.*, 2006).
- The fifteenth top SNP (rs1919550) is a GTEx eQTL for FBXO40 in whole blood, but does not have a high CADD score. However, it lies near a SNP (rs9813391) with a high LLRS that leads to a nonsynonymous change (R145Q) in ARGFX - a homeobox gene - and another SNP (rs4676737) with both a high LLRS and high CADD score (= 14.07) overlapping a repressor region in an intron of FBXO40. The latter SNP is a GTEx eQTL for IQCB1 in fibroblasts, muscular esophagus and thyroid. IQCB1 is associated with Senor-Loken syndrome (Otto *et al.*, 2005), a ciliopathic eye disorder.
- The sixteenth top SNP (rs4955214) is an intronic SNP in OSPBL10.
- The seventeenth top SNP (rs75425359) is one among several high-LLRS SNPs in a cluster upstream of MDGA2, coding for an excitatory synaptic suppressor that has been linked to autism (Bucan *et al.*, 2009; Connor *et al.*, 2016). There are several high-CADD SNPs in this cluster (rs8015664, rs2416146, rs10498399), all of which are intergenic.
- The eighteenth top SNP (rs434923) lies in an intron of PHACTR1 and TBC1D7, and is a GTEx eQTL for both of those genes in different tissues, including nervous, skin, muscular and circulatory. Though not high-CADD-scoring, two neighboring SNPs - rs12525892, rs12525906 - have high CADD scores and high LLRS. rs12525906 is particularly highly conserved across primates, mammals and vertebrates (PhastCons score > 0.97 in all three alignments) and lies in a Segway repressor region within an intron of PHACTR1.
- The nineteenth top SNP (rs144591231) has a high CADD score (= 11.97) and lies in an intergenic region downstream of PUS7L and upstream of IRAK4. PUS7L codes for a protein that catalyzes RNA pseudouridylation (UniProtKB, by similarity), while IRAK4 codes for a kinase involved in the Toll-like receptor pathway (Li *et al.*, 2002), which is an important component of the innate immune response. Deficiency of IRAK4 leads to childhood bacterial and fungal infections (Picard *et al.*, 2003; **?**).
- The twentieth top SNP (rs12147885) is in an intergenic region downstream of SERPINA13P, a pseudogene.
- The twenty-first top SNP (rs73318247) lies in an intron of DOCK2 and is an eQTL of FAM196B - an oncogene (Zhang *et al.*, 2017) - in tibial artery.

- The twenty-second top SNP (rs4946567) is an eQTL of TBC1D32 in cerebellar brain. This SNP has a high CADD score (= 11.02) and is conserved across vertebrates (vertebrate PhyloP = 0.916, vertebrate PhastCons = 0.747). Interestingly, the region in which it is located also harbors signature of selection in Yucatan miniature pigs (Kim *et al.*, 2015; Kwon *et al.*, 2018). TBC1D32 plays a role in cilia assembly (Ko *et al.*, 2010) and may be involved in ciliopathic congenital abnormalities, including midline cleft, microcephaly, and microphthalmia (Adly *et al.*, 2014).

- The twenty-third and twenty-fourth top SNPs (rs5758430, rs4822061) are close to each other and lie in a large region with several high-LLRS SNPs. They are both linked GTEx eQTLs to several genes in a variety of different tissues. They are also both significantly associated with several traits related to body fat, food intake and white blood cells in the UK Biobank GeneATLAS ($P < 10^{-8}$). rs5758430 is associated with lymphocyte percentage and count, eosinophill percentage and count, neutrophill percentage, trunk and body fat percentage, whole body and trunk fat mass, percentage of arm fat in both arms, impedance of both arms and whole body, platelet crit, and intake of tea and processed meat. In turn, rs4822061 is associated with trunk fat percentage and lymphocyte percentage. Although these SNPs do not have particularly high CADD scores, there are several neighboring linked high-LLRS, high-CADD SNPs with significant associations to the same traits: rs2050033 (CADD = 14.73, a regulatory change in a conserved exonic site of MEI1), rs5751148 (CADD = 14.28, an acceptor splice site change in a conserved site of MEI1), rs1807497 (CADD = 12.89), rs132908 (CADD = 12.7), rs739134 (CADD = 10.97, a benign missense mutation of C22orf46), rs5758345 (CADD = 10.58), rs1988153 (CADD = 10.18) and rs5751156 (CADD = 10.15). We also find two significantly-associated SNPs in the GWAS catalog in this region ($P < 10^{-8}$): rs4822024 is associated with Vitiligo (Jin *et al.*, 2012) and rs13054099 is associated with neuroticism (Nagel *et al.*, 2018).

- The twenty-fifth top SNP (rs7978167) is an intronic SNP of TM7SF3, and is an eQTL of MED21 and TM7SF3 in multiple tissues. TM7SF3 is involved in pancreatic insulin secretion (Beck *et al.*, 2011) and cellular stress attenuation (**?**). MED21 codes for a subunit of protein involved in transcriptional activation (Sun *et al.*, 1998).

- The twenty-sixth top SNP (rs76934817) is in an intronic site of XYLT1.

- The twenty-seventh top SNP (rs4629377) lies in the same region as the fifteenth top SNP (rs1919550). It is in an intron of IQCB1 and has a low CADD score (= 1.02).

- The twenty-eight top SNP (rs379236) is an intergenic SNP downstream of SLC37A1. It is an eQTL of RSPH1 in colon and esophagus, of SLC37A1 in skin and of AP001626.2 in tibial nerve.

- The twenty-ninth top SNP (rs4078004) is an intergenic SNP downstream of RBFOX1. There is also a high-LLRS SNP in this region that is significantly associated with insomnia (rs34670506) (Stein *et al.*, 2018).

- The thirtieth top SNP (rs2355805) is an intronic SNP inside AC007682.1.

### Supplementary Tables

Table S1: Top 30 most differentiated SNPs from the European ancestry scan, explicitly looking for highly differentiated loci in the ancestry component that is prevalent among GBR samples. LLRS = log-likelihood ratio score for positive selection.

| chr | pos | rsid | LLRS | CHB | MXL | YRI | GBR | overlapping gene |
|---|---|---|---|---|---|---|---|---|
| 5 | 33951693 | rs16891982 | 22.085902 | 0.0234 | 0.0637 | 0 | 0.9769 | SLC45A2 |
| 15 | 48426484 | rs1426654 | 19.707464 | 0.0306 | 0.1683 | 0.0235 | 1 | SLC24A5 |
| 15 | 28356859 | rs1129038 | 19.290553 | 0 | 0 | 0 | 0.8168 | HERC2 |
| 15 | 28495956 | rs12912427 | 18.270213 | 0.0078 | 0 | 0.0197 | 0.8226 | HERC2 |
| 9 | 16792200 | rs10962596 | 15.819739 | 0 | 0 | 0 | 0.7525 | BNC2 |
| 1 | 1385211 | rs1312568 | 15.066101 | 0.0157 | 0.1245 | 0.0471 | 0.8572 | ATAD3C |
| 2 | 136407479 | rs1446585 | 14.957582 | 0 | 0.0169 | 0 | 0.7398 | R3HDM1 |
| 2 | 136616754 | rs182549 | 14.629386 | 0 | 0 | 0 | 0.7136 | MCM6 |
| 1 | 204784969 | rs3940119 | 14.393216 | 0.1723 | 0.0945 | 0.0158 | 0.8999 | - |
| 4 | 38798648 | rs5743618 | 14.38681 | 0.0234 | 0 | 0.0077 | 0.7131 | TLR1 |
| 2 | 136789259 | rs4954552 | 14.050688 | 0 | 0 | 0 | 0.6999 | - |
| 1 | 1194804 | rs11804831 | 14.042494 | 0.0234 | 0.12 | 0.0236 | 0.8184 | UBE2J2 |
| 2 | 109578855 | rs260687 | 13.981201 | 0.0313 | 0.1385 | 0.1576 | 0.898 | EDAR |
| 2 | 136273578 | rs6735329 | 13.950772 | 0 | 0.0274 | 0 | 0.7176 | ZRANB3 |
| 15 | 91521898 | rs7167128 | 13.917437 | 0 | 0 | 0.0079 | 0.7048 | PRC1 |
| 2 | 136092061 | rs1561277 | 13.893261 | 0 | 0.0199 | 0 | 0.7099 | ZRANB3 |
| 2 | 104832008 | rs11123983 | 13.442969 | 0 | 0 | 0.0627 | 0.7296 | - |
| 10 | 119616558 | rs11198161 | 13.4285 | 0.0548 | 0 | 0.0156 | 0.7311 | - |
| 15 | 52413316 | rs12396 | 13.299209 | 0.0233 | 0.019 | 0.0388 | 0.7356 | GNB5 |
| 12 | 38757712 | rs12423868 | 13.222419 | 0.1715 | 0.0914 | 0.0855 | 0.8893 | - |
| 2 | 135907088 | rs6730157 | 13.21644 | 0 | 0 | 0 | 0.6729 | RAB3GAP1 |
| 17 | 4400392 | rs11657785 | 13.182252 | 0.0078 | 0.2518 | 0 | 0.8486 | - |
| 2 | 104489351 | rs3860446 | 13.157017 | 0.0078 | 0.0029 | 0.0237 | 0.7054 | - |
| 1 | 10767685 | rs284292 | 13.075971 | 0.0702 | 0.0012 | 0.0078 | 0.7366 | CASZ1 |
| 17 | 74809685 | rs72881212 | 13.007157 | 0.0078 | 0 | 0 | 0.6751 | - |
| 14 | 106215480 | rs11624464 | 12.847158 | 0.0154 | 0.0161 | 0 | 0.6967 | - |
| 13 | 111827167 | rs9522149 | 12.845737 | 0.0078 | 0.1275 | 0.039 | 0.7811 | ARHGEF7 |
| 12 | 38884068 | rs10875684 | 12.823057 | 0.1872 | 0.0912 | 0.0933 | 0.8893 | - |
| 16 | 10991539 | rs8057405 | 12.604386 | 0.0705 | 0.0421 | 0.0613 | 0.7816 | CIITA |
| 6 | 145103477 | rs6915047 | 12.601971 | 0.0157 | 0 | 0 | 0.671 | UTRN |

Table S2: Top 30 most differentiated SNPs from the East Asian ancestry scan, explicitly looking for highly differentiated loci in the ancestry component that is prevalent among CHB samples. LLRS = log-likelihood ratio score for positive selection.

| chr | pos | rsid | LLRS | CHB | MXL | YRI | GBR | overlapping gene |
|---|---|---|---|---|---|---|---|---|
| 16 | 48258198 | rs17822931 | 23.271759 | 0.9843 | 0.1223 | 0 | 0.1218 | ABCC11 |
| 16 | 48375777 | rs6500380 | 22.474103 | 0.9922 | 0.1659 | 0 | 0.1038 | LONP2 |
| 1 | 234635790 | rs2175591 | 20.95541 | 0.7056 | 0.0148 | 0 | 0 | - |
| 4 | 100142780 | rs75721934 | 20.453247 | 0.6717 | 0 | 0 | 0 | LOC100507053 |
| 11 | 61579427 | rs72643557 | 20.114033 | 0.6637 | 0 | 0 | 0 | FADS1 |
| 11 | 120154631 | rs12224052 | 19.696284 | 0.6717 | 0.0136 | 0 | 0 | POU2F3 |
| 21 | 43974948 | rs228088 | 19.518001 | 0.8986 | 0.0288 | 0.1012 | 0.1712 | SLC37A1 |
| 11 | 133043841 | rs79802711 | 19.157192 | 0.6406 | 0 | 0 | 0 | OPCML |
| 5 | 128016573 | rs79478220 | 18.476104 | 0.6889 | 0 | 0 | 0.0483 | - |
| 19 | 51441759 | rs11084040 | 18.158963 | 0.8758 | 0.09 | 0.0618 | 0.1292 | - |
| 1 | 169462431 | rs66737090 | 18.123929 | 0.782 | 0.1175 | 0 | 0.0221 | - |
| 4 | 100244319 | rs3811801 | 17.86508 | 0.6092 | 0 | 0 | 0 | - |
| 7 | 5460550 | rs79032618 | 17.863103 | 0.633 | 0 | 0 | 0.0169 | TNRC18 |
| 15 | 28196145 | rs76930569 | 17.232285 | 0.5936 | 0 | 0 | 0 | OCA2 |
| 14 | 32460484 | rs10483393 | 17.091724 | 0.9455 | 0.0448 | 0.3218 | 0.2162 | - |
| 13 | 51047353 | rs9562979 | 16.844984 | 0.7195 | 0.0205 | 0.0308 | 0.0737 | DLEU1 |
| 5 | 127621905 | rs32208 | 16.844522 | 0.6423 | 0 | 0 | 0.0418 | FBN2 |
| 16 | 48864323 | rs4785550 | 16.839722 | 0.7265 | 0.0406 | 0 | 0.0782 | - |
| 20 | 1978301 | rs6136667 | 16.666259 | 0.7502 | 0 | 0 | 0.1521 | - |
| 13 | 74136907 | rs72628045 | 16.34293 | 0.5813 | 0 | 0.0076 | 0 | - |
| 5 | 127908744 | rs12655867 | 16.237287 | 0.6344 | 0 | 0 | 0.0477 | - |
| 14 | 49266305 | rs12589318 | 16.185918 | 0.7811 | 0.1183 | 0.0236 | 0.0569 | - |
| 15 | 64480888 | rs8032157 | 16.109096 | 0.6964 | 0.0679 | 0 | 0.0377 | CSNK1G1 |
| 22 | 42892596 | rs4429562 | 16.103141 | 0.7898 | 0.1445 | 0.0157 | 0.0438 | - |
| 1 | 169176469 | rs1080267 | 16.06272 | 0.8764 | 0.1574 | 0.1634 | 0.0633 | NME7 |
| 1 | 169328403 | rs10800437 | 15.990897 | 0.8764 | 0.1434 | 0.2022 | 0.0629 | NME7 |
| 5 | 58546276 | rs72644097 | 15.965598 | 0.5701 | 0 | 0 | 0.0056 | PDE4D |
| 7 | 5353932 | rs12672490 | 15.868169 | 0.6482 | 0.0248 | 0.0155 | 0.0341 | TNRC18 |
| 2 | 211669698 | rs16844943 | 15.859647 | 0.6096 | 0 | 0.0609 | 0 | - |
| 13 | 50935841 | rs9562961 | 15.846666 | 0.6171 | 0.0364 | 0 | 0.0073 | DLEU1 |

Table S3: Top 30 most differentiated SNPs from the Native American ancestry scan, explicitly looking for highly differentiated loci in the Native American ancestry componet of the MXL samples. LLRS = log-likelihood ratio score for positive selection.

| chr | pos | rsid | LLRS | CHB | MXL | YRI | GBR | overlapping gene |
|---|---|---|---|---|---|---|---|---|
| 14 | 46745012 | rs140736443 | 32.730697 | 0 | 0.6455 | 0 | 0 | LINC00871 |
| 9 | 82968379 | rs6559543 | 27.584847 | 0.0546 | 0.8862 | 0.0234 | 0.117 | - |
| 16 | 80619307 | rs2316155 | 27.399123 | 0.1234 | 0.8028 | 0 | 0 | - |
| 14 | 21647765 | rs77549780 | 27.355769 | 0.0858 | 0.8504 | 0 | 0.0713 | - |
| 12 | 14189549 | rs12425115 | 25.867367 | 0.0078 | 0.7971 | 0.0078 | 0.1331 | - |
| 10 | 8150713 | rs10508343 | 25.609772 | 0.0313 | 0.6238 | 0 | 0 | - |
| 15 | 34936250 | rs16959274 | 25.424824 | 0.1551 | 0.9237 | 0 | 0.09 | - |
| 8 | 4490837 | rs71523639 | 24.59957 | 0.2017 | 0.9116 | 0 | 0.0436 | CSMD1 |
| 1 | 14301862 | rs72640512 | 24.455822 | 0.0703 | 0.8339 | 0 | 0.1191 | - |
| 12 | 29817716 | rs12580697 | 23.967094 | 0.0156 | 0.8022 | 0 | 0.1584 | TMTC1 |
| 14 | 57318110 | rs75607199 | 23.959875 | 0.193 | 0.8766 | 0.0157 | 0.0243 | OTX2-AS1 |
| 2 | 47570021 | rs112850342 | 23.900669 | 0.1162 | 0.786 | 0.031 | 0.028 | LOC101927043 |
| 6 | 155416528 | rs13202771 | 23.654817 | 0.0205 | 0.8396 | 0.1721 | 0.1019 | TIAM2 |
| 4 | 155672155 | rs78441257 | 23.593099 | 0.0078 | 0.625 | 0.0389 | 0.0286 | LRAT |
| 3 | 121364173 | rs1919550 | 23.425263 | 0.068 | 0.7842 | 0 | 0.0871 | HCLS1 |
| 3 | 31907078 | rs4955214 | 23.069388 | 0.0923 | 0.8104 | 0.0148 | 0.085 | OSBPL10 |
| 14 | 48647168 | rs75425359 | 22.793583 | 0.0151 | 0.7649 | 0.0625 | 0.1059 | - |
| 6 | 13283930 | rs434923 | 22.629818 | 0.0156 | 0.6024 | 0 | 0.0427 | PHACTR1 |
| 12 | 44092058 | rs144591231 | 22.400176 | 0.0079 | 0.5264 | 0 | 0.0092 | - |
| 14 | 95117205 | rs12147885 | 22.212869 | 0.0234 | 0.6146 | 0.0156 | 0.0348 | - |
| 5 | 169155152 | rs73318247 | 22.077883 | 0.0623 | 0.7236 | 0.0933 | 0.0284 | DOCK2 |
| 6 | 121674499 | rs4946567 | 21.965283 | 0.093 | 0.7663 | 0 | 0.0793 | - |
| 22 | 42109844 | rs5758430 | 21.933067 | 0.0851 | 0.8876 | 0 | 0.1812 | MEI1 |
| 22 | 42270707 | rs4822061 | 21.583318 | 0.046 | 0.7728 | 0.11 | 0.0719 | SREBF2 |
| 12 | 27161129 | rs7978167 | 21.570297 | 0.0221 | 0.6221 | 0 | 0.0569 | TM7SF3 |
| 16 | 17282105 | rs76934817 | 21.464392 | 0 | 0.6182 | 0 | 0.074 | XYLT1 |
| 3 | 121536106 | rs4629377 | 21.309085 | 0.1001 | 0.7779 | 0 | 0.0791 | IQCB1 |
| 21 | 44012601 | rs379236 | 21.233635 | 0.0392 | 0.9082 | 0.0158 | 0.2599 | - |
| 16 | 7917912 | rs4078004 | 21.153986 | 0.0842 | 0.772 | 0 | 0.0951 | - |
| 2 | 52338125 | rs2355805 | 21.117955 | 0.1158 | 0.7664 | 0 | 0.0649 | - |

Table S4: Top 30 most differentiated SNPs from the scan of the YRI terminal branch. This scan may contain selection candidates specific to the YRI population or selection candidates that are ancestral to the non-African samples, as the ancestry tree used to test for selection is unrooted. LLRS = log-likelihood ratio score for positive selection.

| chr | pos | rsid | LLRS | CHB | MXL | YRI | GBR | overlapping gene |
|---|---|---|---|---|---|---|---|---|
| 8 | 145639681 | rs1871534 | 11.906794 | 0 | 0.0005 | 0.977 | 0 | - |
| 5 | 178626609 | rs6869589 | 11.541667 | 0 | 0.0704 | 0.9844 | 0 | ADAMTS2 |
| 15 | 29427400 | rs10152250 | 11.48232 | 0 | 0.0117 | 0.9923 | 0.0242 | FAM189A1 |
| 1 | 1106112 | rs6670693 | 11.447873 | 0 | 0.0695 | 0.9764 | 0 | - |
| 4 | 3666494 | rs58827274 | 11.341367 | 0 | 0.014 | 0.955 | 0.0085 | - |
| 17 | 2631985 | rs4790359 | 11.118134 | 0 | 0.0446 | 0.9445 | 0 | - |
| 9 | 136769888 | rs2789823 | 11.031687 | 0 | 0.0706 | 0.9552 | 0 | VAV2 |
| 6 | 169656029 | rs6930377 | 10.824098 | 0 | 0.0522 | 0.9446 | 0.0071 | - |
| 17 | 29350769 | rs8073072 | 10.794224 | 0 | 0.0402 | 0.9633 | 0.024 | - |
| 5 | 173642871 | rs10067518 | 10.787147 | 0 | 0 | 0.9166 | 0 | - |
| 1 | 230019048 | rs6587361 | 10.661271 | 0 | 0.0401 | 0.9307 | 0 | - |
| 8 | 145004042 | rs11993782 | 10.657337 | 0 | 0.0062 | 0.9156 | 0 | PLEC |
| 14 | 106215459 | rs989952 | 10.512111 | 0 | 0 | 0.8886 | 0 | - |
| 14 | 57653198 | rs6573130 | 10.454435 | 0 | 0.0571 | 0.945 | 0.0167 | - |
| 9 | 344332 | rs16932430 | 10.409507 | 0 | 0.0568 | 0.9243 | 0 | DOCK8 |
| 4 | 1497319 | rs73794620 | 10.384485 | 0 | 0.0273 | 0.9068 | 0 | - |
| 7 | 145945238 | rs1089605 | 10.295091 | 0.0231 | 0.0573 | 0.9845 | 0.0353 | CNTNAP2 |
| 22 | 39272641 | rs12159761 | 10.285118 | 0 | 0 | 0.8837 | 0 | - |
| 6 | 10661262 | rs7753116 | 10.199686 | 0.0468 | 0.0219 | 0.9386 | 0.0073 | - |
| 9 | 137496040 | rs12004637 | 10.107182 | 0 | 0 | 0.8749 | 0 | - |
| 14 | 105953324 | rs7160340 | 10.070145 | 0 | 0 | 0.8913 | 0.0125 | CRIP1 |
| 2 | 215975232 | rs10180970 | 9.982793 | 0.0234 | 0.0118 | 0.9689 | 0.0503 | ABCA12 |
| 7 | 1092925 | rs73267951 | 9.958967 | 0 | 0 | 0.867 | 0 | C7orf50 |
| 6 | 136494586 | rs6935259 | 9.943145 | 0 | 0.089 | 0.9148 | 0 | PDE7B |
| 8 | 142402629 | rs68107840 | 9.912982 | 0 | 0 | 0.8682 | 0 | - |
| 18 | 67624554 | rs4891384 | 9.900581 | 0 | 0 | 0.9383 | 0.0463 | - |
| 15 | 34271648 | rs592585 | 9.881095 | 0.0078 | 0.0063 | 0.9302 | 0.0359 | AVEN |
| 10 | 111909272 | rs56300906 | 9.797384 | 0.0078 | 0.0417 | 0.9145 | 0.0169 | - |
| 15 | 40759347 | rs679882 | 9.785343 | 0.0468 | 0 | 0.8974 | 0 | BAHD1 |
| 14 | 57833739 | rs2152366 | 9.76709 | 0 | 0.0261 | 0.8669 | 0 | - |

Table S5: [SEPARATE FILE] CADD server annotations (Rentzsch *et al.*, 2018) for top SNPs from Native American ancestry scan that had high log-likelihood ratios in favor of positive selection ($LLRS > 15$) and that were among the top 30 SNPs of the scan or in its surrounding region.
